# Large-scale simultaneous measurement of epitopes and transcriptomes in single cells

**DOI:** 10.1101/113068

**Authors:** Marlon Stoeckius, Christoph Hafemeister, William Stephenson, Brian Houck-Loomis, Pratip K. Chattopadhyay, Harold Swerdlow, Rahul Satija, Peter Smibert

## Abstract

Recent high-throughput single-cell sequencing approaches have been transformative for understanding complex cell populations, but are unable to provide additional phenotypic information, such as protein levels of cell-surface markers. Using oligonucleotide-labeled antibodies, we integrate measurements of cellular proteins and transcriptomes into an efficient, sequencing-based readout of single cells. This method is compatible with existing single-cell sequencing approaches and will readily scale as the throughput of these methods increase.

---

The unbiased and extremely high-throughput nature of modern scRNA-seq approaches has proved invaluable for describing heterogeneous cell populations^1-3^. Prior to the use of single-cell genomics, a wealth of studies achieved detailed definitions of cellular states via cell cytometry by leveraging carefully curated panels of fluorescently labeled antibodies directed at cell surface proteins. Such approaches particularly focused on the immune and nervous systems, where well-characterized markers are reliable indicators of cellular activity and function^4^. Recent studies^5-7^ have demonstrated the potential for coupling ‘index-sorting’ measurements with single-cell transcriptomics, enabling the mapping of immunophenotypes onto transcriptomically-derived clusters. However, recently developed massively parallel approaches based on droplet microfluidics^1-3^, microwells^8,9^ or combinatorial indexing^10,11^ do not utilize cytometry for cellular isolation, and therefore cannot couple protein information with cellular transcriptomes, representing a significant limitation for these exciting approaches.

An ideal approach for cellular phenotyping would combine the measurement of known protein markers with the detection of cellular transcripts in an unbiased manner, while maintaining the high throughput that is obtainable using current scRNA-seq technologies. Recently described attempts to simultaneously detect and/or measure transcripts and proteins in single cells employ proximity ligation or proximity extension assays in combination with digital PCR^12-15^ or mass cytometry^16^. These approaches are limited in scale and/or can only profile a few genes and proteins in parallel (see Supplementary Table 1 for comparison of different technologies).

Here we describe Cellular Indexing of Transcriptomes and Epitopes by sequencing (CITEseq), a method that combines highly multiplexed antibody-based detection of protein markers together with unbiased transcriptome profiling for thousands of single cells in parallel. We demonstrate that the method is readily adaptable to two different high-throughput single-cell RNA sequencing applications and show by example that it can achieve a more detailed characterization of the immune system than scRNA-seq alone. CITE-seq enables analysis of cellular phenotypes at unprecedented detail and readily scales as throughput of scRNA-seq increases.

We hypothesized that a DNA oligonucleotide conjugated to an antibody could be measured by sequencing as a digital readout of protein abundance. We envisioned the oligos to be compatible with oligo dT-based RNA-sequencing library preparations so that they would be captured and sequenced together with mRNAs. To this end, we designed the CITE-seq oligos with the following characteristics: a generic PCR handle for amplification and next-generation sequencing library preparation, a barcode sequence specific for each antibody, and a polyA stretch at the 3’ end (Fig. 1a) designed to anneal to polyT stretches on primers used to initiate reverse transcription. We leverage the DNA-dependent DNA polymerase activity of commonly used reverse transcriptases^17^ (see methods) to convert these barcode-containing CITE-seq oligonucleotides into cDNA during reverse transcription. We adopted a commonly used streptavidin-biotin interaction to link oligos to antibodies^18^ (see methods), and included a disulfide link at the 5’ end of the oligonucleotide, which allows the oligo to be released from the antibody with reducing agents (Supplementary Fig. 1a). The antibody-oligo complexes are incubated with single-cell suspensions using conditions comparable to staining protocols used in flow cytometry^19^. The cells are washed to remove unbound antibodies and are processed for scRNA-seq. In this example, we encapsulated single cells into nanoliter-sized aqueous droplets in a microfluidic apparatus designed to perform Drop-seq^1^ (Fig. 1b). After cell lysis in droplets, both cellular mRNAs and antibody-derived oligos (released in the reducing conditions of the cell lysis buffer – see Supplementary Fig. 1a) anneal to polyT-containing Drop-seq microparticles (Supplementary Fig. 1b) via their 3’ polyA tails. A unique barcode sequence on the oligos attached to the Drop-seq microparticle indexes the cDNA of mRNAs and antibody-oligos of each co-encapsulated cell in the reverse transcription reaction. The amplified antibody-derived tags (hereafter referred to as ADTs) and cDNA molecules can be separated by size and are converted into Illumina-sequencing-ready libraries independently (Supplementary Fig. 1c). Importantly, the two library types are designed to be sequenced together, but because they are generated separately, their relative proportions can be adjusted in a pooled single lane to ensure that the appropriate sequencing depth is obtained for each library.

**Figure 1.**
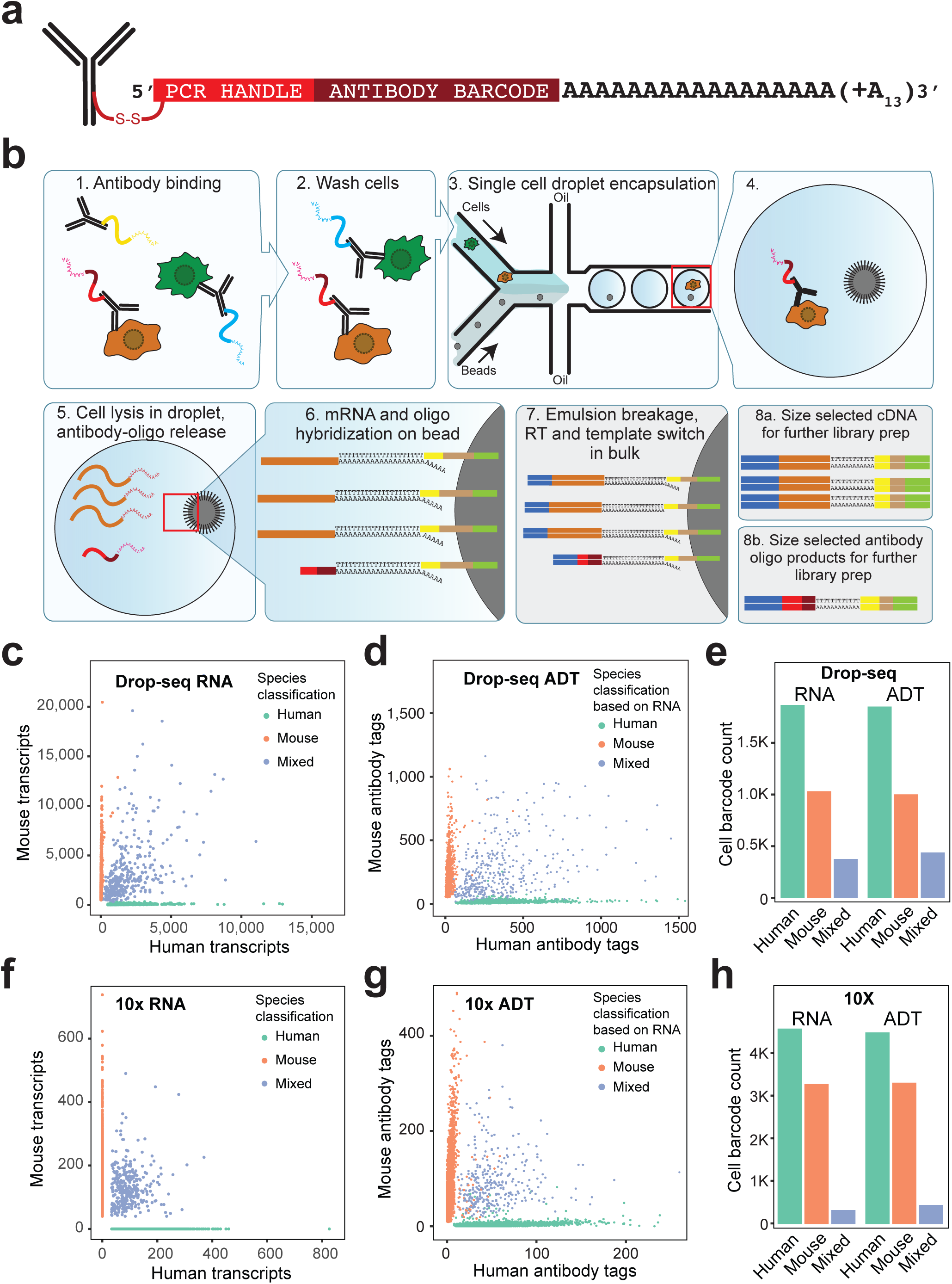
CITE-seq enables simultaneous detection of single cell transcriptomes and protein markers. (**a**) Schematic overview of the DNA-barcoded antibodies used in CITE-seq. Antibodies are linked to oligonucleotides containing an amplification handle, a unique antibody identifier barcode and a polyA tail. A disulfide bond between the antibody and oligo allows release. (**b**) Schematic representation of CITE-seq in combination with Drop-seq^1^. Cells are incubated with antibodies (1). Cells are washed (2) and passed through a microfluidic chip (3) where a single cell and one bead are occasionally encapsulated in the same droplet (4). After cell lysis (5) mRNAs and released antibodyoligos anneal to oligos on Drop-seq beads (6). Reverse transcription and template switch is performed in bulk after emulsion breakage (7). After amplification, full length cDNA and antibody-oligo products can be separated by size and amplified independently (8) (**c - h**) Analysis of mixtures of mouse and human cells that were incubated with oligo-taggedantibodies specific for either human or mouse cell-surface markers (integrin beta CD29) and passed through Drop-seq (c-e) or 10x Genomics (f-h) workflows. In both experiments, we deliberately used a high cell concentration to obtain high rates of doublets (droplets containing two or more cells), to correlate mixed-species transcriptome data to mixed-species protein signals from individual droplets. (**c** and **f**) quantification of the number of human and mouse transcripts associating to each cell barcode. Each green point indicates a cell barcode (i.e. droplet containing one or more cells) from which we measured >90% human transcripts; each red point indicates a cell barcode with >90% mouse transcripts. Blue points indicate cell barcodes (i.e. droplets) from which we observed a mixture of human and mouse transcripts. (**d** and **g**) Quantitation of species-specific antibody tags (ADTs) associated with each cell barcode. Points are colored based on species classifications using transcripts in (c) and (f), respectively. (**e** and **h**) Quantification of human, mouse or mixed-cell barcodes based on RNA transcripts, or ADTs, detected in Drop-seq and 10x Genomics workflows.

To assess the ability of our method to distinguish single cells based on surface protein expression, we designed a proof-of-principle ‘species-mixing’ experiment. A mixed suspension of human (HeLa) and mouse (4T1) cells was incubated with a mixture of DNA-barcoded anti-mouse and anti-human CD29 (Integrin beta-1) antibodies. After washing to remove unbound antibodies, we performed droplet-based single-cell analysis (Drop-seq^1^) to investigate the concordance between species of origin of the transcripts and ADTs (Fig. 1c-e, Supplementary Fig. 2a). 97.3% of droplets that were identified as containing either human cells (green) or mouse cells (red) by transcriptome alignment (Fig. 1c), were also identified as having surface-bound epitopes corresponding to the appropriate species (Fig. 1d). Similarly, 99.4% of droplets that were first identified by the antibody-based approach to contain either human or mouse cells showed the same result by transcriptome analysis. In parallel, we tested the performance of the method in a commercially available system from 10x Genomics and obtained comparable results (Fig. 1fh, Supplementary Fig. 2b). In both experiments, we deliberately used a high cell concentration to obtain high rates of doublets (droplets containing two or more cells), to demonstrate that we can detect more than one ADT type per droplet, and to correlate mixed-species transcriptome data with mixed-species ADT signals from individual droplets. 96% (Drop-seq) and 89% (10x Genomics) of droplets that contained human and mouse cell mixtures (Fig. 1c and f, blue) also had ADTs from both human and mouse antibodies (Fig. 1d and g, blue). Cell counts based on RNA or ADT are comparable between both methods (Fig. 1e and h). The consistency between the species classification of cell barcodes based on detected transcripts and ADTs demonstrates the specificity of this method.

We next wanted to charaterize the quantitative nature of our CITE-seq ADT protein readout. Flow cytometry is the gold-standard for identification and enumeration of cell subsets based on quantitative differences in surface markers, particulary within the immune system^20,21^. We therefore first aimed to benchmark the sensitivity of CITE-seq protein detection to flow cytometry using immune cells as a model system. For this we prepared a set of CITE-seq antibodies directed against markers commonly used in flow cytometry to identify select major immune populations (e.g. CD3 and CD19) and to discriminate relevant immune sub-populations (e.g. CD4 and CD8). We then, in parallel, performed multiparameter flow cytometry (Fig. 2a) and CITE-seq (Fig. 2b) experiments using the same set of antibodies on aliquots of a single pool of peripheral blood mononuclear cells. Using ADT levels we were able to construct cytometry-like ‘bi-axial’ gating plots (Fig. 2b) and compare these qualitatively and quantitatively to the visualized flow cytometry data (Fig. 2a). Cell distribution profiles and expression patterns for proteins associated with T cells (CD3, CD4, CD8, CD57, CD2) and B cells (CD19) were remarkably similar (Fig. 2a,b), as were those for plasmacytoid and myeloid dendritic cells and monocytes (CD11c, CD14; Supplmental Fig. 3a). Moreover, the percentages of cells identified by flow cytometry and CITE-seq within the different immune subsets were also highly similar (Supplemental Fig. 3b).

**Figure 2.**
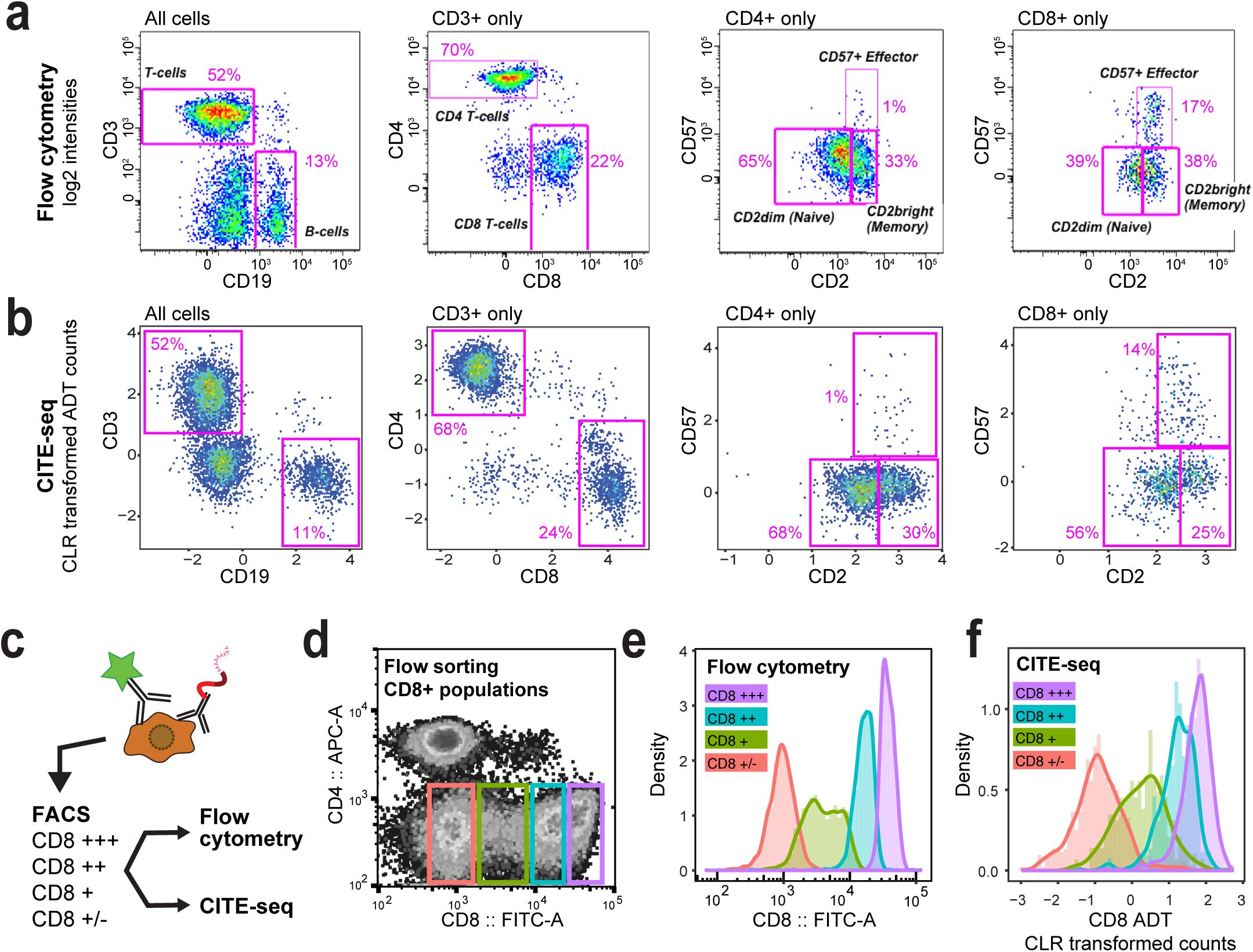
Qualitative and quantitative comparison between CITE-seq and flow cytometry. **(a-b)** Comparison of qualitative readout of flow cytometry to CITE-seq. Cells from the same aliquot were processed for flow cytometry (a) and CITE-seq (b). Functionally important immune subsets were selected based on their established flow cytometry expression patterns and their relative frequencies compared to the entire population, and within the CD3, CD4 and CD8 positive subsets. **(c)** Schematic overview of the experiment performed to compare quantitative readout in CITE-seq to flow cytometry. Cord blood mononuclear cells were stained with a mixture of monoclonal antibodies tagged either with the fluorophore FITC, or with a DNA barcode targeting CD8a. Cells were then sorted into four different bins of CD8 expression levels by FACS. Each sorted pool was divided into two fractions which were measured by flow cytometry or CITE-seq. (**d**) Profile of CD4 and CD8 fluorescence in CBMCs. Colored boxes are gates set to sort different levels of CD8 expression. **(e)** Flow cytometry of cells sorted in panel d. Merged histograms of CD8 levels in the four different pools of cells. **(f)** CD8 levels of the different pools of cells sorted in panel d, as measured by CITE-seq. Merged histograms of four CITE-seq runs of the separate pools.

Next, we asked whether quantitative differences in marker expression, as observed by flow cytometry, can be recapitulated by CITE-seq. For this, we used the CD8 protein as a model, since it is expressed at a wide range of levels across immune cell populations including cytotoxic T cells, natural killer cells, and dendritic cells. We incubated cord blood mononuclear cells (CBMCs) with CITE-seq antibody conjugates and fluorophore-conjugated antibodies, so that some CD8 epitopes on each cell would be labeled by fluorophore and some by oligo. Cells were sorted (by fluorescence-activated cell sorting, FACS) into separate pools based on CD8 fluorescence (CD8 very high (+++), CD8 high (++), CD8 intermediate(+) and CD8 low(+/−); Fig. 2c,d, Supplementary Fig. 3c). Each pool was then split and separately reanalyzed by flow cytometry and CITE-seq. For each pool defined by FACS, similar expression levels of CD8 were observed by both methods (compare Fig. 2 e and f, Suppementary Fig. 3d and e); moreover, we also observed the expected quantitative differences in CD3, CD16 and CD56 ADT levels across pools. For example, CD3 is most highly expressed in the cells with the highest CD8 signal, as expected for cytotoxic T cells (Supplementary Fig. 3d). We also observed higher CD56 and CD16 levels in the pools of cells expressing lower levels of CD8, as expected for NK cells (see below, Supplementary Fig. 3f). We conclude that CITE-seq ADT levels are compatible with immunophenotyping, yielding results consistent with gold-standard flow cytometry.

We next aimed to perform a broad immunophenotypic and transcriptomic characterization of a complex immune cell population using CITE-seq. The immune system has been extensively profiled by both cell surface markers^20^ and scRNAseq ^3,6,22^, which allows us to identify celltypes based on surface markers and transcriptome, and therefore is an ideal system to validate the multi modal readout of CITE-seq. We prepared a CITE-seq panel of 13 well-characterized monoclonal antibodies that recognize specific cell-surface proteins routinely used as markers for immune-cell classification, including CD3, CD4, CD8, CD56, CD19, CD11c, CD16, CD14, CD45RA and CD34 (Supplementary Table 2). Measuring protein abundance by antibodies can be complicated by non-specific binding, resulting in higher background and/or false positive results. To estimate background of antibody binding within experiments we capitalized on the multimodal readout obtained from CITE-seq. We reasoned that cells from a different species would be 1) unlikely (or in many cases, demonstrated not) to have epitopes that specifically cross-react with our antibodies and 2) be easily distinguished transcriptomically. We therefore spiked a low number of mouse fibroblasts (~4% 3T3) into our CBMCs, incubated the cell pool with our immune CITE-seq antibody panel and ran the 10x Genomics single cell workflow. We measured both cDNA and ADT profiles from 8,005 cells (see methods). Unsupervised graph-based clustering based on RNA expression resulted in partitioning of the identified cell barcodes into recognizable cell types indicated by expression of select marker genes (Fig. 3a, Supplementary Fig. 4), corresponding broadly to T cells, B cells, NK cells, erythrocytes, precursors and monocytes (Supplementary Fig. 4). We first asked whether our detected ADT levels were consistent with our unbiased transcriptomic clustering. We observed very low levels of ADT counts in mouse cells which allowed us to set a baseline for signal vs noise to more clearly delineate positive from negative cell populations (Supplementary Fig. 5a,b). Through this thresholding step, we identified three antibody-oligo conjugates with no specific binding (*i.e.*, no signal over background threshold) and excluded these from further analysis (Supplementary Fig. 5b).

**Figure 3.**
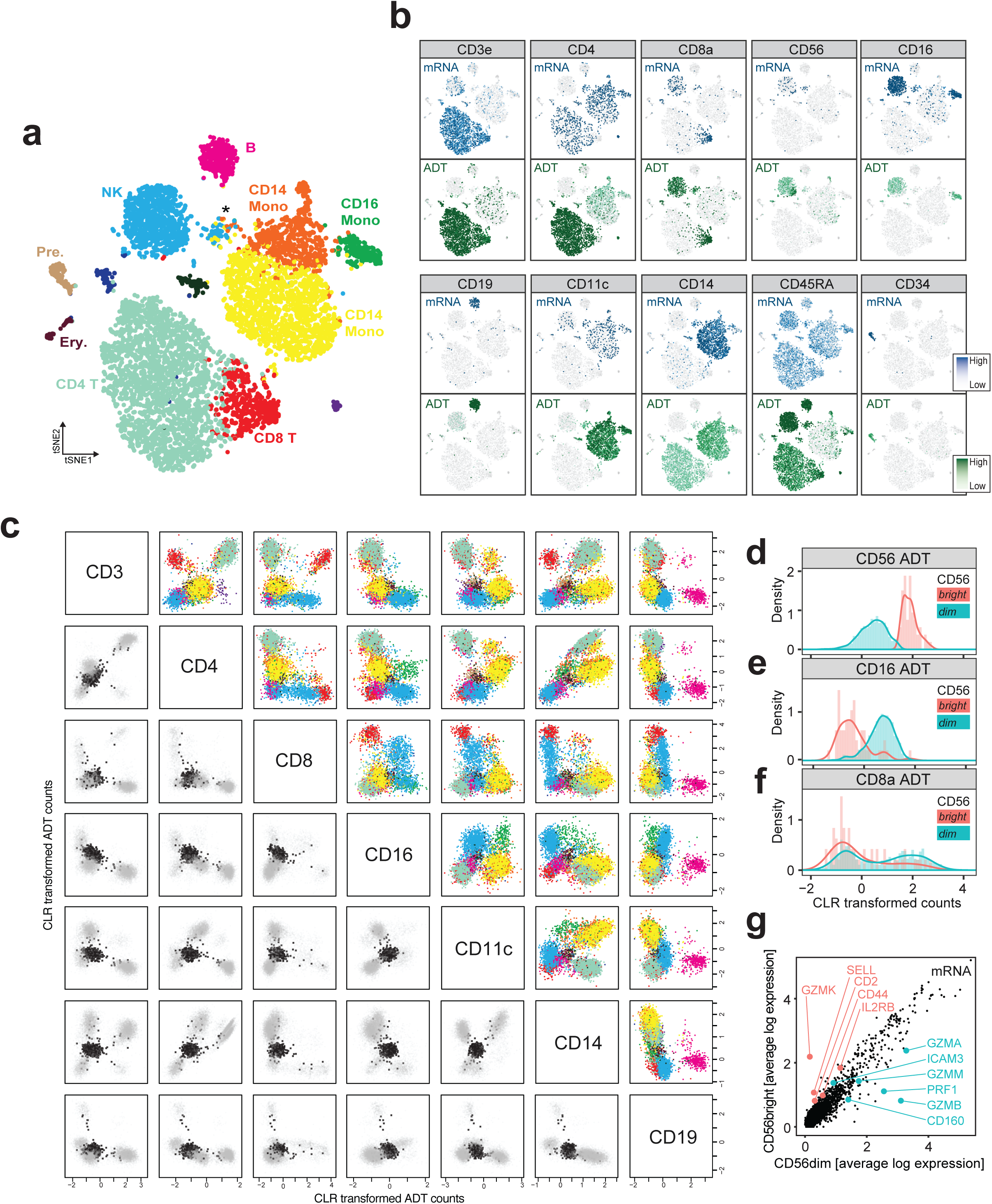
CITE-seq allows detailed classification of cord blood mononuclear cells. (**a**) Clustering of 8,005 CITE-seq single-cell expression profiles of CBMCs reveals distinct cell populations based on transcriptome. The plot shows a two-dimensional representation (tSNE) of global gene expression relationships among all cells (methods). Major cell types in cord blood can be discerned based on marker gene expression (Supplementary Fig. 4). CD4 and CD8 T cells, B cells, NK cells, CD16 monocytes, CD14 monocytes, precursors, erythrocytes, and putative doublets co-expressing multiple lineage markers (*) are indicated. The mouse control cell population was excluded from the clustering. (**b**) mRNA (blue) and corresponding ADT (green) signal for the CITE-seq antibody panel projected on the tSNE plot from (a). Darker shading corresponds to higher levels measured. (**c**) Multimodal bi-axial plots. Pairwise comparison of different ADT levels in single cells for select markers (see Supplementary Fig. 5c for all markers). Upper right: ADT counts in CBMCs, colors based on RNA clusters shown in panel a. Lower left: ADT counts from mouse control cells (black) that were spiked in at very low frequency are overlaid on the CBMC data (light grey). ADT counts were centered log-ratio (CLR) transformed (see methods). (**d-f**) Analysis of NK cells. NK cells were split into CD56 bright and dim groups based on CD56 ADT levels. Histogram of CD56 (d), CD16 (e), CD8a (f) levels in the CD56^dim^ and CD56^bright^ groups. (**g**) Differential gene expression analysis between the CD56^bright^ and CD56^dim^ cells. Genes known from literature to be higher expressed in CD56^bright^ are marked in red, genes known to be higher in CD56^dim^ are marked in green.

For surface markers that are known to be exclusive to individual immune populations, we detected ADT enrichment in the correct populations: We observe the T cell marker CD3 within the T cell population (Fig. 3b), and CD4 and CD8 in largely non-overlapping T cell subpopulations (Fig. 3b). We observe CD19 ADTs almost exclusively in the B cell cluster (Fig. 3b), CD56 and CD8 ADTs in the NK cluster, and CD11c, CD14 in the monocyte and dendritic cell cluster, together with CD16, which is also observed in the NK cells (Fig. 3b). Importantly, we can also correctly identify a very rare precursor cell population present at less than 2% in cord blood (CD34+ cells; Fig 3b). We observed that the ADT levels per-cell exhibited higher counts than mRNA-levels of the same genes, likely due to the presence of greater copy number of surface proteins than mRNA demonstrating that ADTs were less prone to ‘drop-out’ events. Consistent with this, we find low correlations between mRNA and ADT on a single cell basis and higher correlation when averaging expression within clusters (Supplementary Fig. 6). Spiking-in mouse cells at low frequency also allowed us to estimate the true doublet rates (4%) in our experiments, and compare these to the estimates provided by the equipment manufacturer (~3.1%). We then used the ADT levels and clustering information based on transcriptome to construct multimodal CITE-seq ‘bi-axial’ gating plots (Fig. 3c). These revealed very similar profiles and the same relationships that are well-established by flow cytometry, including strong anti-correlation of CD4 and CD8 ADT levels indicative of mutually exclusive expression. Since we are able to distinguish the celltypes based on RNA within the plots, we were able to resolve quantitative differences in marker expression between subsets, such as expression differences of CD8 between NK and T cells (blue and red cells; Fig. 3c) or CD4 between monocytes and T cells (yellow and turquoise cells; Fig. 3c). In addition to clustering cells based on their transcriptome, we performed unbiased clustering exclusively based on ADT levels, identifying clear and consistent cell type separation (Supplementary Fig. 7).

We next asked whether CITE-seq could enhance our characterization of immune cell phenotypes, compared to scRNA-seq data alone. We noted an opposing gradient of CD56 and CD16 ADT levels within our transcriptomically-derived NK cell cluster, potentially corresponding to CD56^bright^ and CD56^dim^ subsets^23,24^ (Fig. 3b). We therefore sub-divided our NK cell cluster based on CD56 ADT levels (Fig. 3d). Consistent with known literature, we observed an apparent complementarity between levels of CD16, and to a lesser extend of CD8, to CD56 ADTs between these two subsets (Fig. 3e,f)^23,24^. Analyzing differentially expressed transcripts between the two sub-clusters allowed us to validate 10 out of 11 known marker genes for which we could detect expression^23-25^ (including GZMB, GZMK and PRF1) as differentially expressed between these two cell types (Fig. 3g). We emphasize that the distinction between CD56^bright^ and CD56^dim^ has not previously been observed in scRNA-seq analysis, even on massive datasets consisting of >250,000 cells ^3^, illustrating the potential for integrated and multi-modal analyses to enhance our discovery and description of cellular phenotypes, particularly when differentiating between cell populations with subtle transcriptomic differences.

The single-cell community is increasingly interested in deriving more than one layer of information from single cells^26^. CITE-seq enables multi-modal analysis of single cells at the scale afforded by droplet-based single-cell sequencing approaches. This will not only provide a more detailed characterization of cell populations, but also allow the study of post-transcriptional (and post-translational) gene regulation in single cells at an unprecedented depth. We demonstrate the value of multi-modal analysis to reveal phenotypes that could not be discovered by using scRNA-seq alone. In particular, joint analysis of protein and RNA levels enables us to finely resolve NK cell (CD56^bright^ vs CD56^dim^) populations that have only subtle transcriptomic differences. CITE-seq is readily scalable in terms of both the number of cells analyzed, and the number of protein markers interrogated. Detection of oligo-barcoded antibodies is not limited by signal collision; a modest 10 nucleotide sequence can easily encode more barcodes than there are human proteins. In contrast, flow cytometry, which relies on the separation of emission spectra of fluorophores to detect different signals, only allows for the routine measurement of 4-6 parameters per cell, or up to 30 parameters in highly specialized labs^27,28^. The high multiplexing potential of CITE-seq also far exceeds mass cytometry-based parallelization (CyTOF up to 100 tags^28^), and we have shown that it maintains a comparable quantitative readout to flow cytometry. In addition, we envision that mild cell permeabilization and fixation procedures that are used for intracellular antibody staining in signaling-specific cytometry assays^28^ will also be compatible with CITE-seq, which may significantly expand the number of markers and biological questions that can be interrogated by this approach. A modified version of CITE-seq in which only ADTs are analyzed on a massively parallel scale without any attempt to capture cellular mRNAs (essentially cytometry by sequencing) can also be envisaged, and a conceptually similar approach, Abseq, has very recently been described^29^. In contrast to CITE-seq, Abseq requires highly advanced custom microfluidics and multiple microfluidic devices to perform multiple operations in droplets, and only allows for the detection of protein markers.

Finally, we have shown that the method works on a commercially-available single-cell instrument (10x Genomics) and should be readily adaptable to other droplet, microwell and combinatorial indexing-based high-throughput single-cell sequencing technologies^2,8-11^ with no, or minor customizations. We believe that CITE-seq has the potential to advance single-cell biology by layering an extra dimension on top of single-cell transcriptome data.

## Methods

### Conjugation of Antibodies to DNA-barcoding oligonucleotides

Highly specific, flow cytometry tested monoclonal antibodies (see below) were conjugated to oligonucleotides containing unique antibody identifier sequences and a polyA tail. Oligonucleotides were conjugated to antibodies by streptavidin-biotin conjugation using the LYNX Rapid Streptavidin Antibody Conjugation Kit (Bio-Rad, USA), according to manufacturer’s instructions with modifications. Specifically, we labeled 15μg of antibody with 10μg of streptavidin. At this ratio, an average of two streptavidin tetramers will be conjugated per antibody molecule, which results in 4-8 binding sites for biotin on each antibody. DNA-oligonucleotides were purchased (IDT, USA) with a 5’ biotin modification or with a 5’ amine modification and biotinylated using NHS-chemistry according to manufacturer’s instructions (EZ Biotin S-S NHS, Thermo Fisher Scientific, USA). The disulfide bond allows separation of the oligo from the antibody with reducing agents. Separation of the oligo from the antibody may not be needed for all applications. Excess Biotin-NHS was removed by gel filtration (Micro Biospin 6, Bio-Rad) and ethanol precipitation. Streptavidin-labeled antibodies were incubated with biotinylated oligonucleotides in excess (1.5x theoretically available free streptavidin) overnight at 4°C in PBS containing 0.5M NaCl and 0.02% Tween. Unbound oligo was removed from antibodies using centrifugal filters with a 100KDa MW cutoff (Millipore, USA). Removal of excess oligo was verified by 4% agarose gel electrophoresis (Supplementary Fig. 1a). Antibody-oligo conjugates were stored at 4°C supplemented with sodium azide and BSA. See supplementary protocol for a more detailed description.

### List of Antibodies used for CITE-seq

See Supplementary Table 2 for list antibodies, clones and barcodes used for CITE-seq.

### Antibody-oligo sequences

We leverage the DNA-dependent DNA polymerase activity of commonly used reverse transcriptases^17^ to convert CITE-seq oligonucleotides into cDNA during reverse transcription together with mRNAs. The DNA-dependent DNA polymerase activity of MMLV reverse transcriptases is well established. All SMART (Switching Mechanism at 5’ end of RNA Template) library prep protocols (e.g. commercialized by Clontech) rely on this activity: The RT enzyme switches at the end of the RNA template to a template switch oligo (TSO), which is mainly DNA, for further cDNA synthesis. This activity has been shown to be highly sensitive and reproducible. Single cell RNA-seq protocols (including 10x Genomics and Drop-seq) also entirely rely on this activity to append a PCR handle to the 5’end of full length cDNAs which is used for subsequent amplification. Depending on the application the PCR-amplification handle in the antibody-barcoding oligos must be changed depending on which sequence read is used for RNA readout (*e.g.* 10x Single Cell 3’v1 uses read 1 while Drop-seq and 10x Single Cell 3’ v2 use read 2). Our proof-of-principle human/mouse antibody-barcoding oligonucleotide designs included UMIs, which are redundant for Drop-seq and 10x protocols due to the UMI addition to the cDNA at reverse transcription. UMIs on the antibody-conjugated oligonucleotide may be useful for other iterations of the method where UMIs are not part of the scRNA-seq library preparation protocol.

Species mixing – Drop-seq (containing Nextera read2 handle)

~~~
BC6:  /5AmMC12/GTCTCGTGGGCTCGGAGATGTGTATAAGAGACAGGCCAATNNBAAAAAAAAAAAAAAAAAAAAAAAAAAAAAAAAAAA
BC12: /5AmMC12/GTCTCGTGGGCTCGGAGATGTGTATAAGAGACAGCTTGTANNBAAAAAAAAAAAAAAAAAAAAAAAAAAAAAAAAAAA
~~~

Species mixing – 10x (Single cell 3’ version 1, Nextera read1 handle)

~~~
BC6:  /5AmMC12/TCGTCGGCAGCGTCAGATGTGTATAAGAGACAGGCCAATNNBAAAAAAAAAAAAAAAAAAAAAAAAAAAAAAAAAAA
BC12: /5AmMC12/TCGTCGGCAGCGTCAGATGTGTATAAGAGACAGCTTGTANNBAAAAAAAAAAAAAAAAAAAAAAAAAAAAAAAAAAA
~~~

CBMC profiling – (Drop-seq and 10x v2 compatible oligos, containing TruSeq small RNA read 2 handle)

~~~
v2_BC1:  /5AmMC12/CCTTGGCACCCGAGAATTCCAATCACGBAAAAAAAAAAAAAAAAAAAAAAAAAAAAAAAA
v2_BC2:  /5AmMC12/CCTTGGCACCCGAGAATTCCACGATGTBAAAAAAAAAAAAAAAAAAAAAAAAAAAAAAAA
v2_BC3:  /5AmMC12/CCTTGGCACCCGAGAATTCCATTAGGCBAAAAAAAAAAAAAAAAAAAAAAAAAAAAAAAA
v2_BC4:  /5AmMC12/CCTTGGCACCCGAGAATTCCATGACCABAAAAAAAAAAAAAAAAAAAAAAAAAAAAAAAA
v2_BC6:  /5AmMC12/CCTTGGCACCCGAGAATTCCAGCCAATBAAAAAAAAAAAAAAAAAAAAAAAAAAAAAAAA
v2_BC9:  /5AmMC12/CCTTGGCACCCGAGAATTCCAGATCAGBAAAAAAAAAAAAAAAAAAAAAAAAAAAAAAAA
v2_BC10: /5AmMC12/CCTTGGCACCCGAGAATTCCATAGCTTBAAAAAAAAAAAAAAAAAAAAAAAAAAAAAAAA
v2_BC12: /5AmMC12/CCTTGGCACCCGAGAATTCCACTTGTABAAAAAAAAAAAAAAAAAAAAAAAAAAAAAAAA
v2_BC8:  /5AmMC12/CCTTGGCACCCGAGAATTCCAACTTGABAAAAAAAAAAAAAAAAAAAAAAAAAAAAAAAA
v2_BC11: /5AmMC12/CCTTGGCACCCGAGAATTCCAGGCTACBAAAAAAAAAAAAAAAAAAAAAAAAAAAAAAAA
v2_BC13: /5AmMC12/CCTTGGCACCCGAGAATTCCAAGTCAABAAAAAAAAAAAAAAAAAAAAAAAAAAAAAAAA
v2_BC14: /5AmMC12/CCTTGGCACCCGAGAATTCCAAGTTCCBAAAAAAAAAAAAAAAAAAAAAAAAAAAAAAAA
v2_BC5:  /5AmMC12/CCTTGGCACCCGAGAATTCCAACAGTGBAAAAAAAAAAAAAAAAAAAAAAAAAAAAAAAA
~~~

### Cell DNA-barcoded antibody labelling for CITE-seq

Cells were resuspended in cold PBS containing 2% BSA and 0.01% Tween and filtered through 40μm cell strainers (Falcon, USA) to remove potential clumps and large particles. Cells were then incubated for 10 minutes with Fc receptor block (TruStain FcX, BioLegend, USA) to block non-specific antibody binding. Subsequently cells were incubated in with mixtures of barcoded antibodies for 30 minutes at 4°C. Antibody concentrations were 1μg per test as recommended by the manufacturer (Biolegend, USA) for flow cytometry applications. Cells were washed 3x by resuspension in PBS containing 2% BSA and 0.01% Tween, followed by centrifugation (~480x*g* 5 minutes at 4°C) and supernatant exchange. After the final wash cells were resuspended at appropriate cell concentration in PBS for Drop-seq^1^ or 10x Genomics^3^ applications

### Drop-seq – CITE-seq

Drop-seq was performed as described^1^ with modifications. For the human/mouse mixing experiment cells were loaded at a concentration of 400 cells/μL to achieve a high doublet rate. For PMBC experiments cells were loaded at 150 cells/μL. cDNA was amplified for 10 cycles and products were then size separated with Ampure Beads (Beckman Coulter, USA) into <300 nt fragments containing antibody derived tags (ADTs) and >300 nt fragments containing cDNAs derived from cellular mRNA. ADTs were amplified 10 additional cycles using specific primers that append P5 and P7 sequences for clustering on Illumina flowcells. Alternatively, antibody tags can be amplified directly from thoroughly washed Drop-seq beads after RNA-cDNA amplification using specific primers for the antibody oligo and Drop-seq bead-RT oligo. cDNAs derived from RNA were converted into sequencing libraries by tagmentation as described^1^. After quantification libraries were merged at appropriate concentrations (10% of a lane for ADT, 90% cDNA library). Sequencing was performed on a HiSeq 2500 Rapid Run with v2 chemistry per manufacturer’s instructions (Illumina, USA).

### 10x – CITE-seq

The 10x single cell run was performed according to the manufacturer’s instructions (10x Genomics, USA) with modifications. For the Human/Mouse mixing experiment (Single Cell 3’ v1) 10,000 cells were loaded to yield an intermediate/high doublet rate. For CBMC profiling (Single Cell 3’v2), 4,000 cells were loaded to yield a doublet rate <5%. cDNA was amplified for 10 cycles and products were then size separated with Ampure Beads (Beckman Coulter, USA) into <300 nt fragments containing antibody derived tags (ADTs) and >300 nt fragments containing cDNAs derived from cellular mRNA. ADTs were amplified 10 additional cycles using specific primers that append P5 and P7 sequences for clustering on Illumina flowcells. A sequencing library from cDNAs derived from RNA was generated using a tagmentation based approach akin to that used in Drop-seq for the Single Cell 3’ v1 experiments, or according to manufacturer’s instructions for the Single Cell 3’v2 experiments. ADT and cDNA libraries were merged and sequenced as described above.

### Cell culture

HeLa (human), 4T1 (mouse) and 3T3 (mouse) cells were maintained according to standard procedures in Dulbecco’s Modified Eagle’s Medium (Thermo Fisher, USA) supplemented with 10% fetal bovine serum (Thermo Fisher, USA) at 37°C with 5% CO2. For the species mixing experiment, HeLa and 4T1 cells were mixed in equal proportions and incubated with DNA barcoded CITE-seq antibodies as described above. For the low frequency mouse spike-ins ~5% 3T3 cells were mixed into CBMC pool before performing CITE-seq.

### Blood mononuclear cells

Cord blood mononuclear cells (CBMCs) were isolated from cord blood (New York Blood Center) as described^30^. Cells were kept on ice during and after isolation. Peripheral blood mononuclear cells were obtained from Allcells (USA).

### Comparing FACS and CITE-seq

Cells were stained with a mixture of fluorophore (FITC) labelled antibodies and CITE-seq oligo labelled antibodies from the same monoclonal antibody clone (RPA-T8) targeting CD8a, at concentrations recommended by the manufacturer (1ug per test, Biolegend, USA). Cells were also stained with Anti-CD4-APC antibody (RPA-T4, Biolegend, USA). Cells were sorted into pools of different CD8 expression levels using the Sony SH800 cell sorter, operated per manufacturer’s instructions. Pools were then split into two and reanalyzed by flow cytometry using Sony SH800 and processed for CITE-seq using Drop-seq as described above. Flow cytometry data was plotted using FlowJo (USA).

### Multiparameter flow cytometry

Cells were stained with the following mouse anti-human antibodies, purchased from BD Biosciences (San Jose, CA): CD3 Hilyte 750 Allophycocyanin (H7APC), CD4 Brilliant Blue(BB) 630, CD8a Phycoerythrin (PE), CD14 Brilliant Violet (BV) 750, CD19 BV570, CD11c Cyanin5 PE, CD2 Brilliant Ultraviolet (BUV) 805, and CD57 BB790. After washing cells in PBS and fixing in 0.5% paraformaldehyde, samples were acquired on a BD Symphony A5 flow cytometer and data was analyzed using FlowJo v9 (Eugene, OR).

## Computational methods

### Single cell RNA data processing and filtering

The raw Drop-seq data was processed with the standard pipeline (Drop-seq tools version 1.12 from McCarroll lab). 10x data from the species mixing experiment was processed using Cell Ranger 1.2 using default parameters and no further filtering was applied. 10x data from CBMC experiments (v2 chemistry) was processed using the same pipeline as Drop-seq data. Reads were aligned to the human reference sequence GRCh37/hg19 (CD8 FACS comparison), or an hg 19 and mouse reference mm10 concatenation (species mixing experiment, CBMCs).

Drop-seq data of the species mixing experiment was filtered to contain only cells with at least 500 UMIs mapping to human genes, or 500 UMIs mapping to mouse genes. For the CD8 FACS comparison data, we kept only cells with PCT_USABLE_BASES >= 0.5 (fraction of bases mapping to mRNA, this is part of the metrics outputted by the default processing pipeline). We further removed any cells with less than 200 genes detected and cells with a total number of UMIs or genes (in log10 after adding a pseudo-count) that is more than 3 standard deviations above or below the mean. The same filtering strategy was used for the CBMC data, the only difference being a gene threshold of 500.

### Single cell ADT data processing and filtering

Antibody and cell barcodes were directly extracted from the reads in the fastq files. Since the antibody barcodes were sufficiently different in the species mixing experiment, we also counted sequences with Hamming distance less than 4. For the CBMCs we counted sequences with Hamming distance less than 2. Reads with the same combination of cellular, molecular and antibody barcode were only counted once.

We kept only cells that passed the RNA-specific filters and had a minimum number of total ADT counts (species mixing: 10, CD8 FACS comparison: 1, CBMC: 50).

### CBMC RNA normalization and clustering

After read-alignment and cell filtering, we assigned the species to each cell barcode. If more than 90% of UMI counts were coming from human genes, the cell barcode was considered to be human. If it was less than 10% the assigned species was mouse. Cell barcodes in between were considered mixed species. The resulting assignment was human: 8005, mouse: 579, mixed: 33. Unless stated otherwise, analysis was performed on only the human cells and genes from the human reference genome.

We converted the matrix of UMI counts into a log-normalized expression matrix *x* with 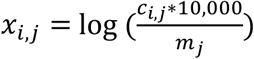, where *c_i,j_* is the molecule count of gene *i* in cell *j* and *m_j_* is the sum of all molecule counts for cell *j*. After normalization each gene was scaled to have mean expression 0 and variance 1.

We identified 556 highly variable genes by fitting a smooth line (LOESS, span=0.33, degree=2) to log10(var(UMIs)/mean(UMIs)) as a function of log10(mean(UMIs)) and keeping all genes with a standardized residual above 1 and a detection rate of at least 1%.

To cluster the cells, we performed dimensionality reduction followed by modularity optimization. We ran principal component analysis (PCA) using the expression matrix of variable genes. To determine the number of significant dimensions, we looked at the percent change in successive eigenvalues. The last eigenvalue to feature a reduction of at least 5% constituted our significant number of dimensions (in this case the number was 13). For clustering we used a modularity optimization algorithm that finds community structure in the data^31^. The data is represented as a weighted network with cells being nodes and squared Jaccard similarities as edge weights (based on Euclidian distance of significant PCs and a neighborhood size of 40 (0.5% of all cells)). The clustering algorithm, as implemented in the “cluster_louvain” function of the igraph R package, find a partitioning of the cells with high density within communities as compared to between communities. For 2D visualization we further reduced the dimensionality of the data to 2 using t-SNE^32,33^.

### CBMC ADT normalization and clustering

Since each ADT count for a given cell can be interpreted as part of a whole (all ADT counts assigned to that cell), and there are only 13 components in this experiment, we treated this data type as compositional data and applied the centered logratio (CLR) transformation (Aitchison 1989). Explicitly, we generated a new CLR-transformed ADT vector *y* for each cell where 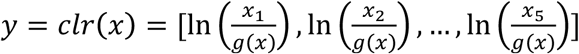 and *x* is the vector of ADT counts including one pseudocount for each component, and *g*(*x*) is the geometric mean of *x*. We noticed that the ADT counts were on slightly different scales for the different antibodies, perhaps due to differences in antibody specificity and/or epitope abundance. To compensate for the resulting shifts in the non-specific baseline ADT signal, we examined the density distribution of the CLR-transformed ADT counts of all antibodies separately for human and mouse cells (Supplementary Fig. 5a,b). For each ADT we determined the mean and variance of the mouse cells and defined the species-independent cutoff (separating ‘off’ state from ‘on’ state where protein is present) to be one standard deviation larger than the mean.

To cluster cells based on ADT counts, the same general approach as for the RNA data was taken, except no dimensionality reduction was performed. Instead we subtracted the mouse-derived cutoffs from the CLR-transformed ADT counts for each antibody. Cell-to-cell weights were squared Jaccard similarities based on Euclidean distance and neighborhood size of 0.5% of the total number of cells.

### Estimation of doublet rate using low frequency mouse spike-in

For estimation of the doublet rate in our experiments we modeled the droplet cell capture process as a Poisson distribution with a loading rate lambda and a fixed mouse fraction of 6.5%. We optimized lambda so that simulated data would most closely match the observed species distribution. The resulting lambda was 0.068 and the doublet rate (fraction of droplets with more than one cell of all droplets with at least one cell) observed in the simulations was 4%.

## Acknowledgements

We acknowledge members of the NYGC Technology Innovation lab S. Jaini and K. Pandit for critical discussions and support. We thank E. Papalexi for help with CBMCs isolation. We thank M. Coppo, S. Fennessey, B. Baysa and S. Pescatore at NYGC for sequencing support. We thank C. Kocks for discussions of the manuscript. Multiparameter flow cytometer instrument time and reagents were generously provided by the Vaccine Research Center of the National Institutes of Health. CH was supported by a Deutsche Forschungsgemeinschaft research fellowship.

---

**Supplementary Figure 1.**
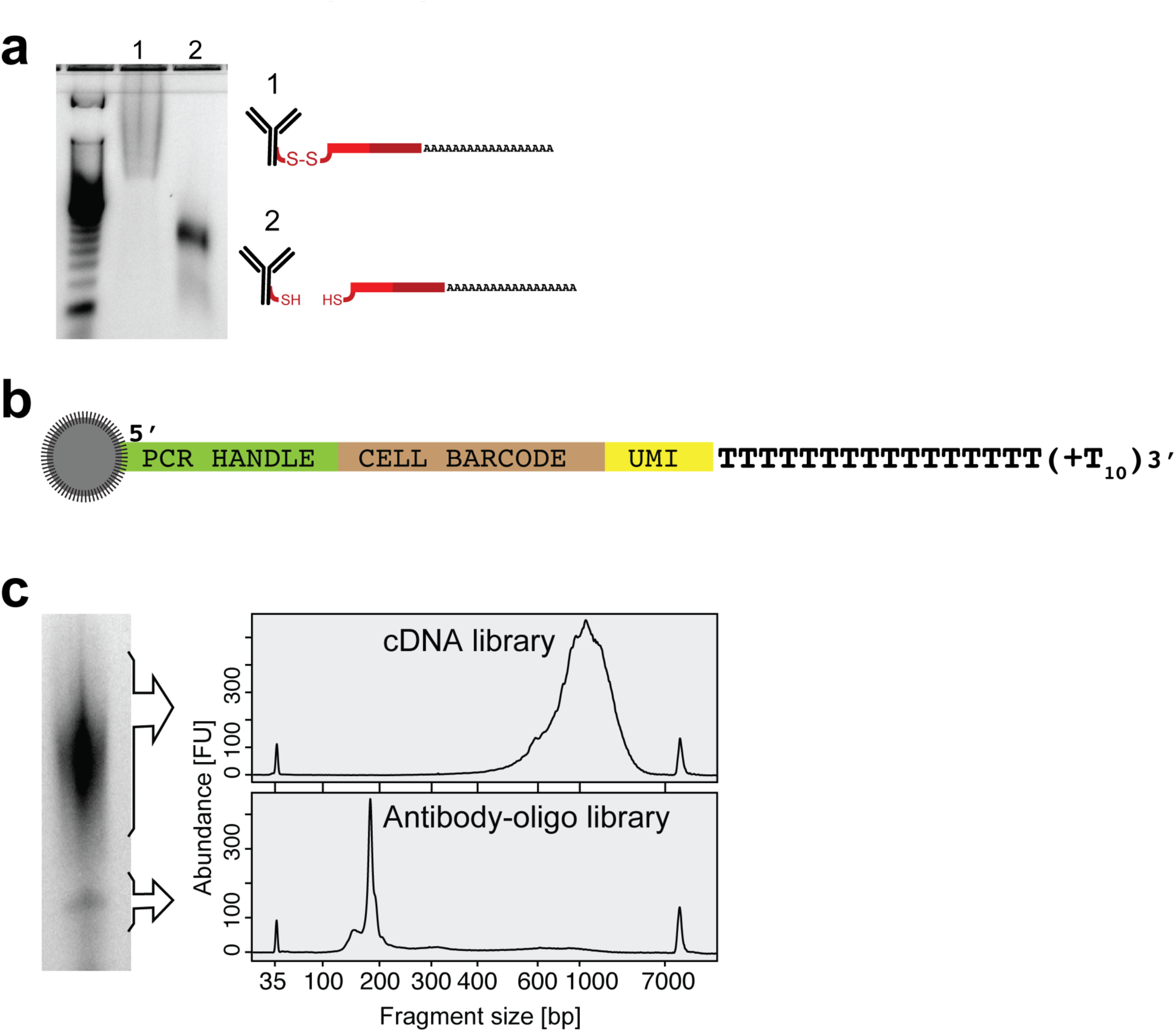
CITE-seq library preparation. (**a**) Antibody-oligonucleotide complexes appear as a high-molecular-weight smear when run on an agarose gel (1). Cleavage of the oligo from the antibody by reduction of the disulfide bond collapses the smear to oligo length (2). (**b**) Drop-seq beads are microparticles containing oligonucleotides with a common PCR handle, a unique cell barcode, followed by a unique molecular identifier (UMI) and a polyT tail^1^. (**c**) Reverse transcription and amplification produces two product populations with distinct sizes (left panel). These can be size separated and amplified independently to obtain full length cDNAs (top panel, capillary electrophoresis trace) and ADTs (bottom panel, capillary electrophoresis trace).

**Supplementary Figure 2.**
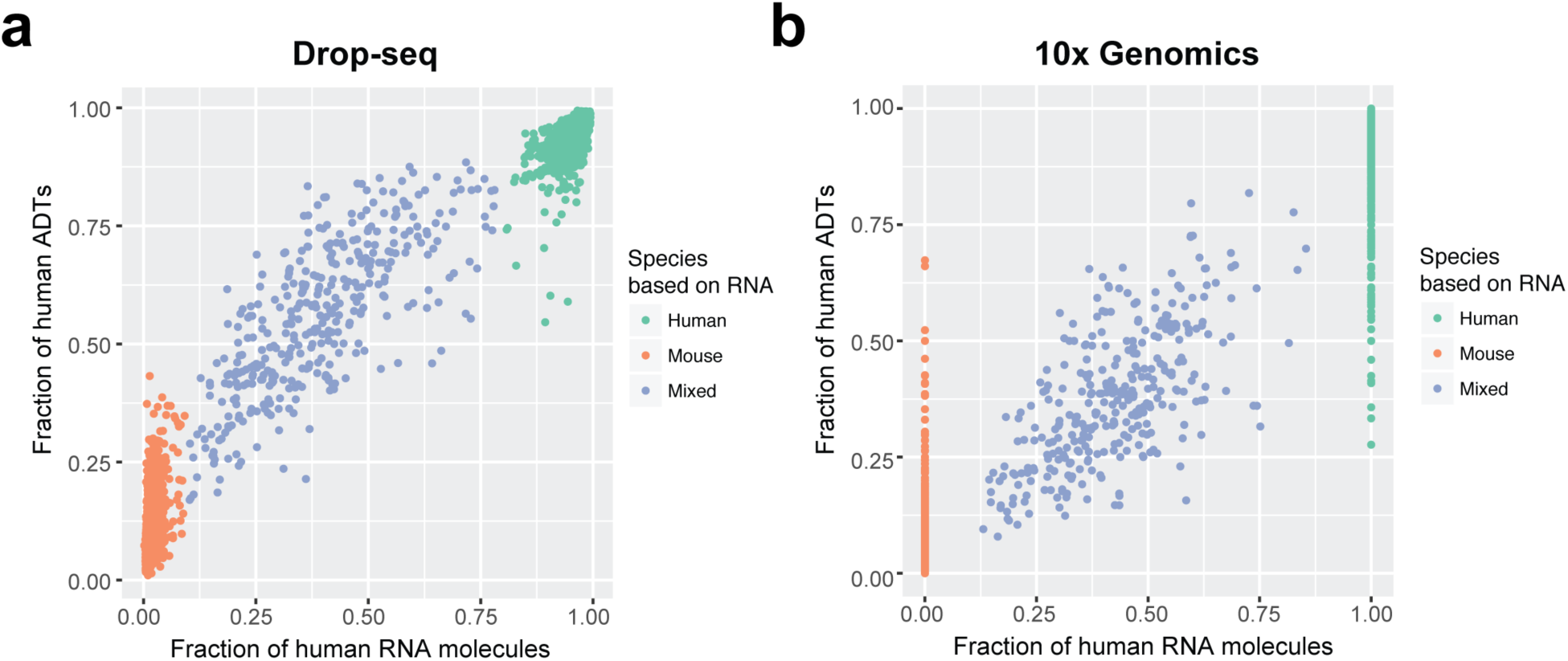
Human and mouse RNA and ADT fractions. **(a-b)** Analysis of mixtures of mouse (4TI) and human (HeLa) cells that were incubated with oligo-tagged-antibodies specific for human or mouse cell-surface markers (integrin beta, CD29) and passed through the Drop-seq (**a**) or 10x Genomics (**b**) workflow. Each point represents one cell barcode (*i.e.*, droplet containing one or more cells). Fractions of human RNA molecules compared to human ADTs for detected cell barcodes.

**Supplementary Figure 3.**
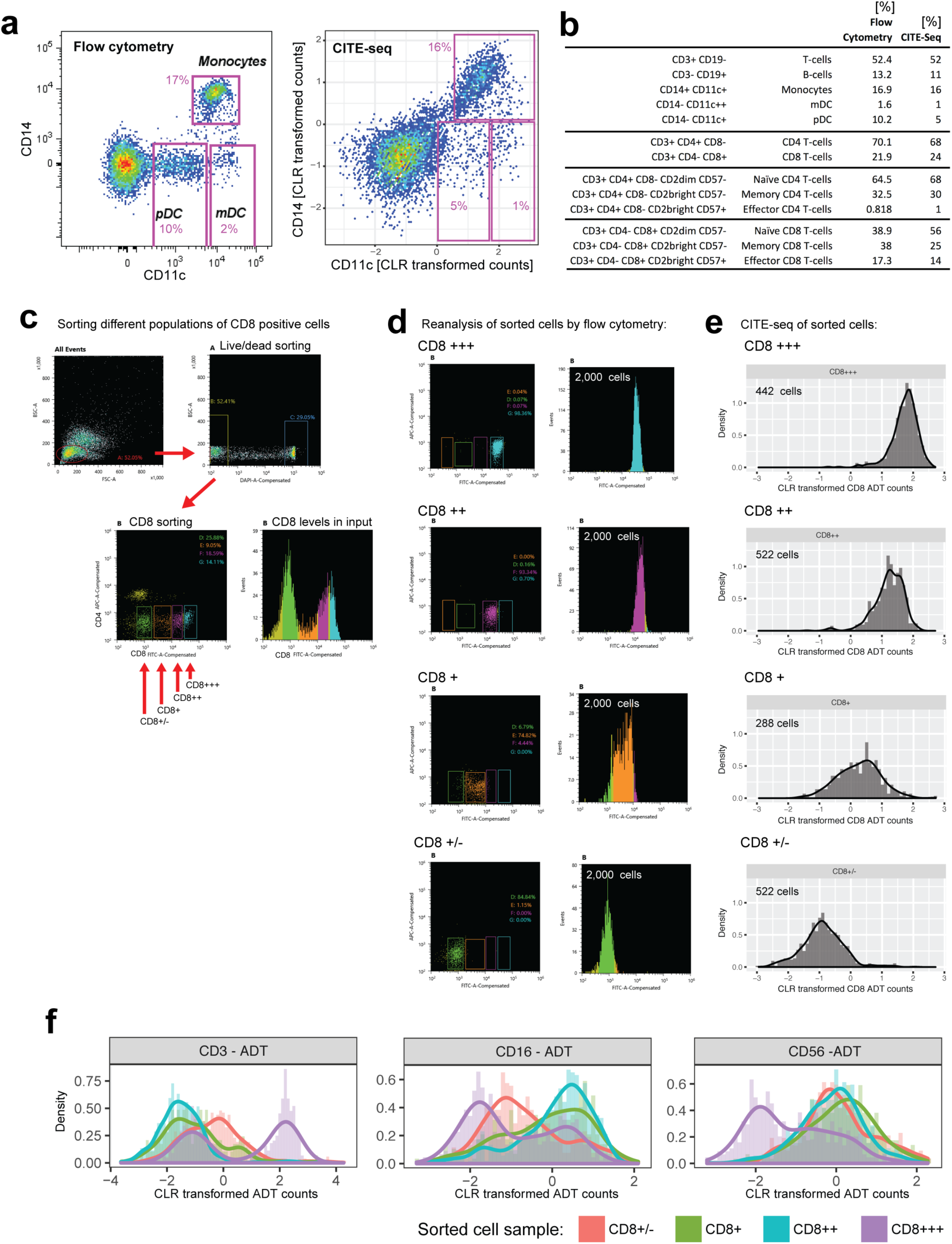
Comparing qualitative and quantitative readout in CITEseq and flow cytometry. **(a-b)** Periperal blood mononuclear cells were processed by flow cytometry and CITE-seq to compare qualitative readout between both technologies. Relevant immune populations were labelled (a) and their abundances relative to the entire population compared (b, see also Fig. 2a,b). (**b**) Relative abundances of relevant immune cell subsets as determined by flow cytometry and CITE-seq (see Fig. 2a,b; and panel a) **(c)** Cord blood mononuclear cells were stained with a mixture of monoclonal antibodies targeting CD8a tagged with the fluorophore FITC and with a DNA barcode. Cells were then passed through a FACS and sorted into four different bins of CD8 expression levels. Each sorted pool was then divided into two fractions and passed once more through a flow cytometer and processed for CITE-seq. (**c**) Cells were first gated based on forward and sideward scatter and separated from dead cells. Profile of CD4 and CD8 fluorescence in CBMCs. Colored boxes are gates set to sort different levels of CD8. (**d**) Re-analysis of cells sorted into CD3 very-high (+++), high (++), intermediate (+) and low (+/−) by flow cytometry. Histograms of CD8 levels (fluorescence intensities) in the four different pools of cells. 2,000 cells were measured for each run. **(e)** CD8 levels obtained by CITE-seq of the different pools of pools of cells sorted in panel a. Histograms of four CITE-seq runs of the separate pools. 288-522 cells were measured for each run. **(f)** Levels of CD3, CD14 and CD56 as determined by CITE-seq in the different CD8 pools. Merged histograms of the four CITE-seq runs.

**Supplementary Figure 4.**
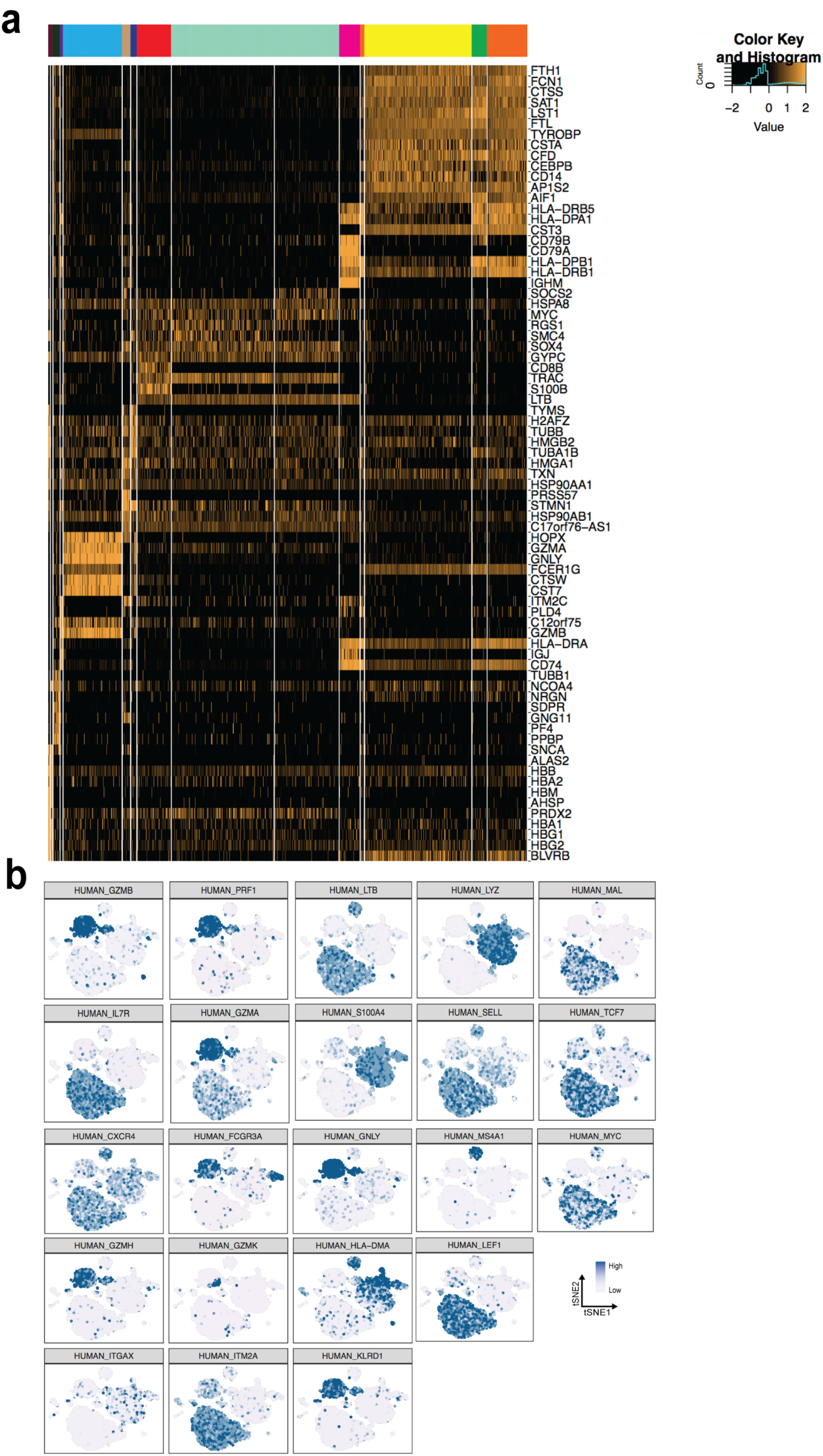
Marker gene expression in CBMC clusters. (**a**) Gene expression heatmap of most differentially expressed marker genes used to define clusters. Dimensionality reduction followed by modularity optimization was used to cluster 8,005 CBMCs (methods). Cluster color assignments are identical to Figure 3a. The mouse control cell population was excluded from the clustering. (**b**) Expression of individual marker genes in the context of tSNE representation of cell relationships based on single-cell gene-expression profiles. Levels of transcripts corresponding to specific marker genes are indicated by blue shading. Plots were used to define cell types in different clusters.

**Supplementary Figure 5.**
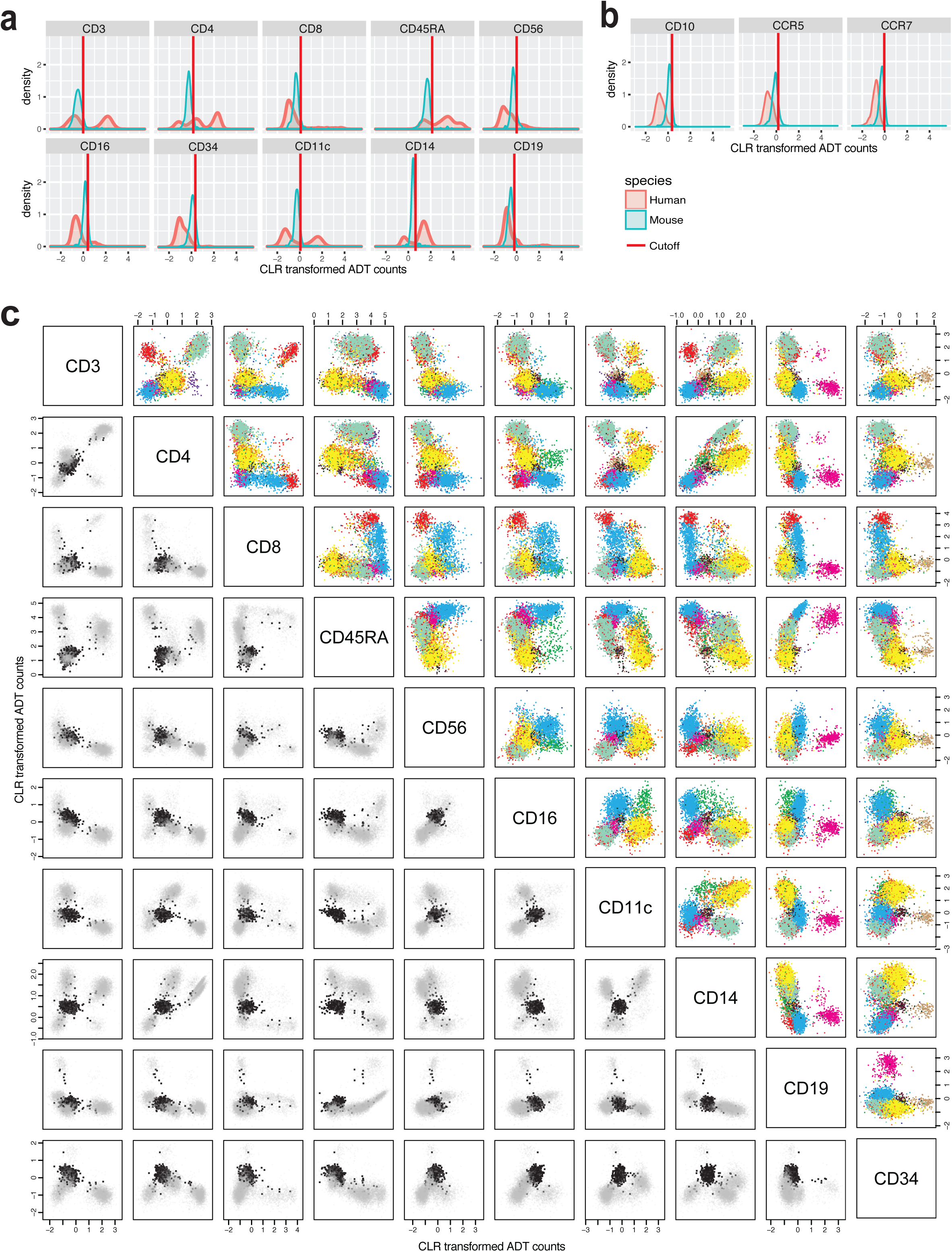
Resolving single-cell RNA clusters using ADTs. **a)** Histograms of CLR-transformed ADT counts in CBMCs (red) and mouse control cells spiked at very low frequency (blue). Solid line shows the determined cutoff for significant ADT signal (mean of mouse values + standard deviation of mouse values, see methods). (**b**) Histograms of CLR-transformed ADT counts for the three antibodies-oligo conjugates in our 13 antibody pool that did not pass the mouse-derived threshold. (**c**) Multimodal biaxial plots. Pairwise comparison of different CLR-transformed ADT levels in CBMCs. Upper right: colors based on RNA clusters shown in Figure 3a. Lower left: mouse control cells (black) that were spiked at very low frequency are overlaid the CBMCs (light grey).

**Supplementary Figure 6.**
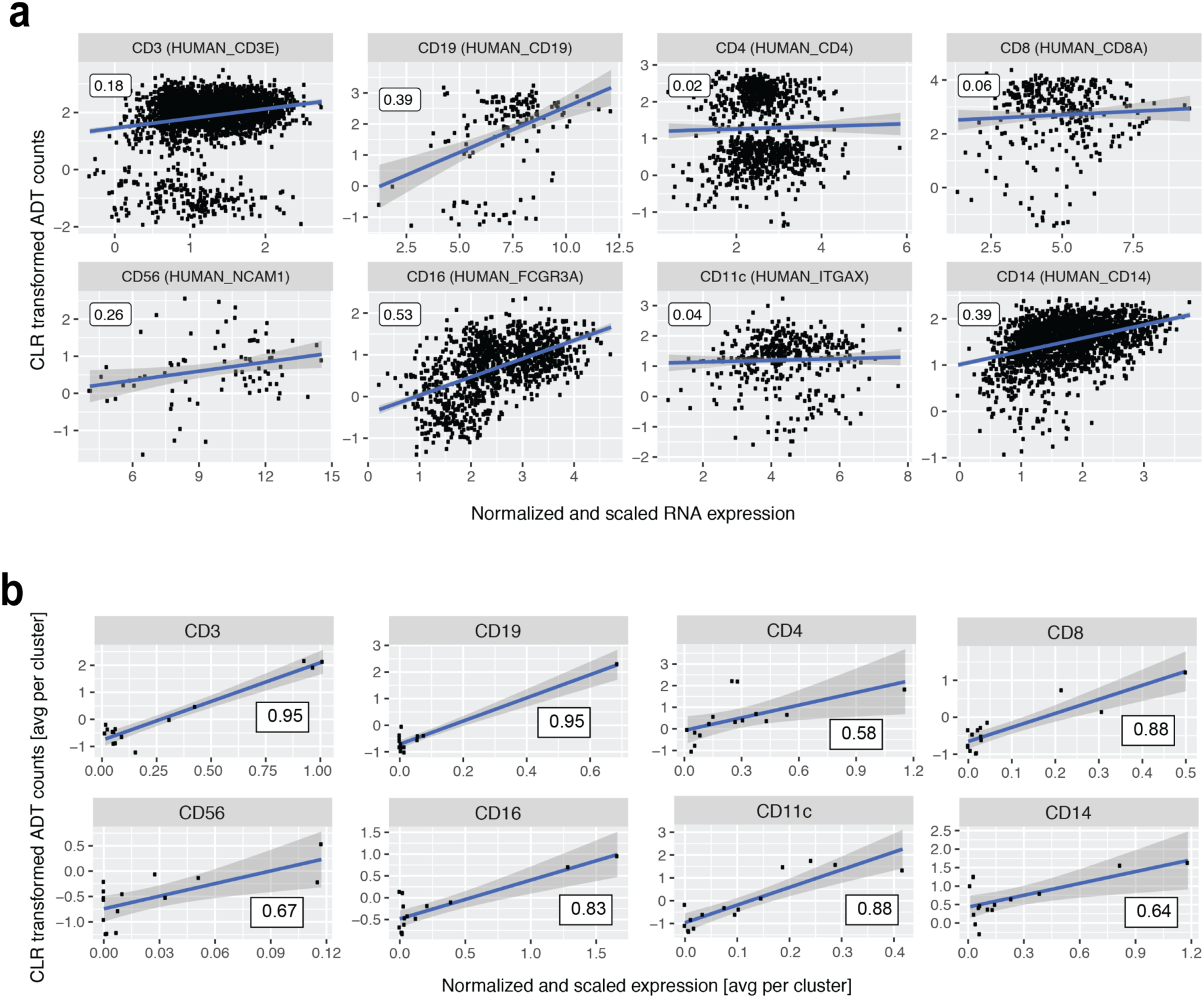
Correlations between mRNA and protein marker levels in CITE-seq. (**a**) Correlation between normalized mRNA expression and CLR-transformed ADT counts at a single cell level. For each gene, cells with no detected mRNA molecules were excluded. Pearson’s correlation coefficient is shown in the boxed labels. (**b**) Correlation between normalized mRNA expression and CLR-transformed ADT counts at the cluster level. No cells were excluded and mRNA and ADT signals were averaged per cluster before calculating Pearson’s correlation coefficient.

**Supplementary Figure 7.**
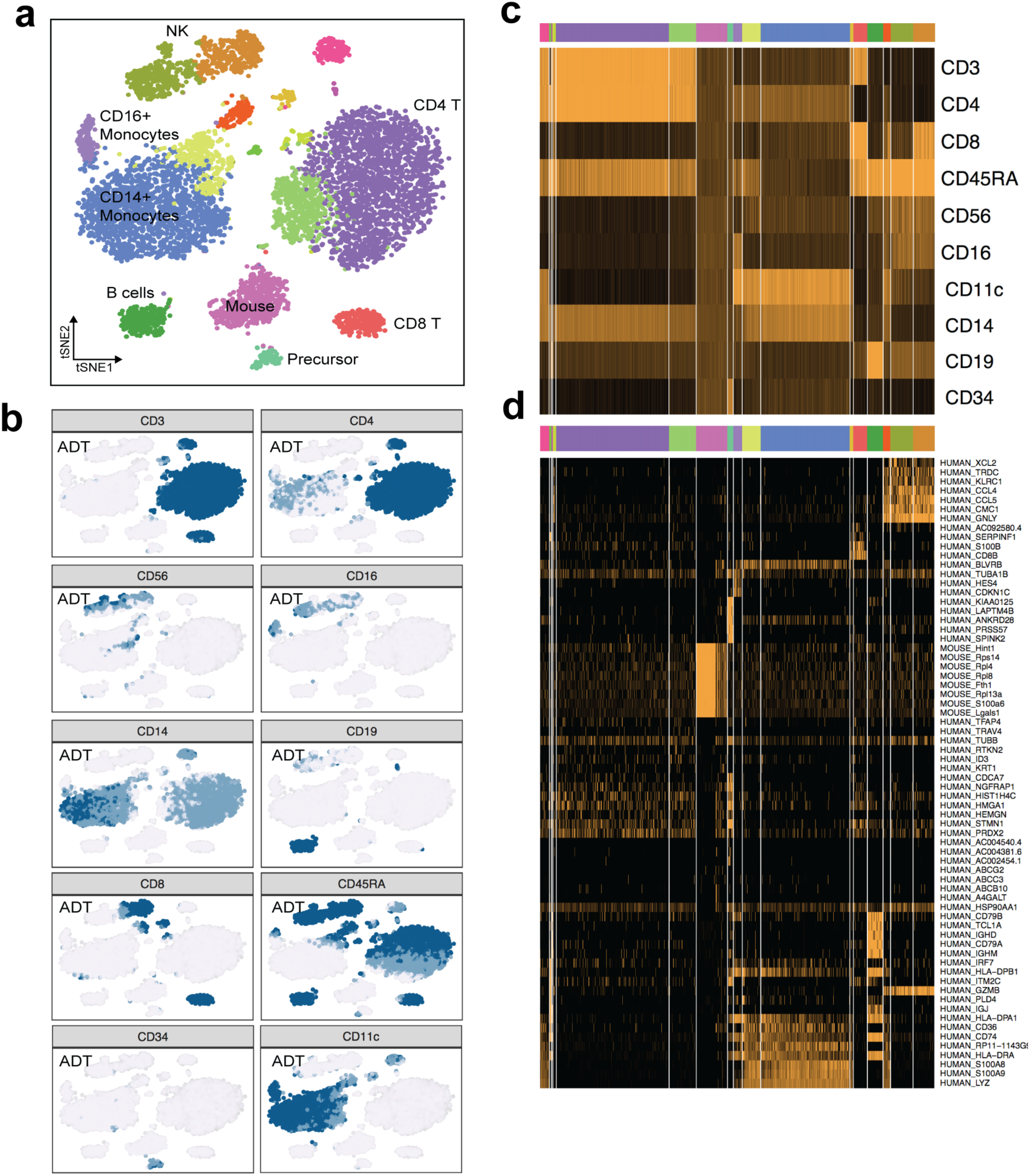
Clustering of CBMCs based on ADTs. (**a**) CITE-seq single-cell ADT profiles of ~8,005 CBMCs (and ~600 mouse control cells) were clustered using modularity optimization resulting in 17 cell populations (including the mouse control cell population) with distinct antibody compositions. (**b**) ADT levels for 10 markers in clusters defined by ADT levels. Levels of ADT are indicated by blue to dark blue shading. (**c**) Single-cell ADT level heatmap in the ADT-derived clusters. Colors of clusters in top panel represent colors in panel a. (**d**) Single-cell gene-expression heatmap of top marker genes of ADT-derived clusters. Colors of clusters in top panel represent colors in (a).

**Supplementary Table 1.** Overview of methods that allow multiplexed measurements of RNA and/or proteins in single cells.

| Technology | # transcripts | # proteins | # cells | Reference |
| --- | --- | --- | --- | --- |
| Single-cell RNA-seq (droplet based) | > 4,000 | - | > 10,000 | 1, 2, 3 |
| CyTOF | - | < 100 | > 10,000 | 28 |
| FACS | - | < 30 | > 10,000 | 27 |
| Index-sorting RNA-seq | > 4,000 | < 30 | ~ 96 - hundreds | 5, 6, 7 |
| Wafergen ICELL8 | > 4,000 | ~ 4 | ~ 1,800 |  |
| Fluidigm C1 | > 4,000 | ~ 4 | ~ 96 - ~ 800 |  |
| PLA/PEA & qPCR | ~ 96 | ~ 36 | ~ 96 - hundreds | 12, 13, 14, 15 |
| PLAYR | ~ 20 | ~ 20 | > 10,000 | 16 |

**Supplementary Table 2.**
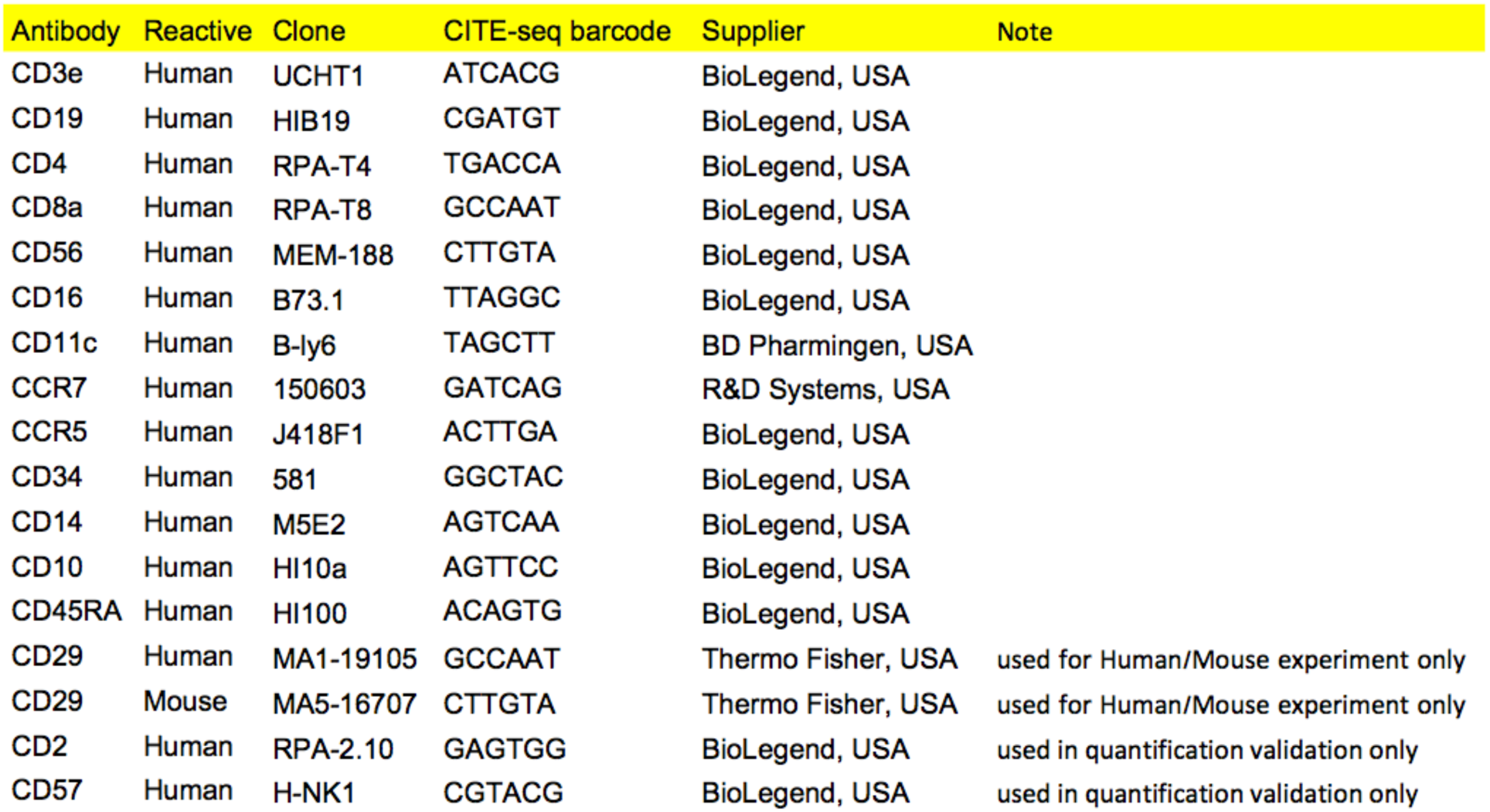
List of antibody clones, supplier and barcodes.

